# “Bolder” together – response to human social cues in free-ranging dogs

**DOI:** 10.1101/760108

**Authors:** Debottam Bhattacharjee, Shubhra Sau, Anindita Bhadra

**Author notes:** Correspondence Behaviour and Ecology Lab, Department of Biological Sciences, Indian Institute of Science Education and Research Kolkata, Mohanpur Campus, Mohanpur, PIN 741246, West Bengal, INDIA., (AB).

## Abstract

Interspecific interactions within an ecosystem have different direct and indirect effects on the two interacting species. In the urban environment, humans are a part of an interaction network of several species. While indirect human influence on different urban species has been measured extensively, experimental studies concerning direct human influence are lacking. In this study, we tested interactions between groups of urban free-ranging dogs (*Canis lupus familiaris*) and solitary unfamiliar humans in ecologically relevant contexts. We provided different sets of dogs with four commonly used human social cues (neutral, friendly, low and high impact threatening) to understand their responses at the group-level and identify potential inter-individual differences. Finally, we compared data from a previous study to investigate the differences in behavioural outcomes between solitary and groups of dogs while interacting with humans. The study not only strengthens the idea of situation-relevant responsiveness in free-ranging dogs but also highlights the minute differences between solitary and group-level reactions in the form of higher approach and less anxious behaviour of groups towards the unfamiliar human. Additionally, we report inter-individual differences and the effect of sex while responding to the threatening cues. Our study suggests a direct benefit of group-living over a solitary lifestyle in free-ranging dogs while interacting with humans in the streets.

**Summary statement:** Free-ranging dogs can benefit by living in groups over a solitary lifestyle while interacting with unfamiliar humans in urban habitats irrespective of having significant inter-individual differences.

## Introduction

Behavioural adjustments during interspecific interactions are widespread in the animal world. Such interactions can involve both positive and negative stimuli from either or both the individuals of the interacting species. Of particular interest is how humans directly or indirectly influence the behaviour and personality of other animals living close to them. A range of species has been shown to alter their behaviour upon indirect human influence, especially in the context of urbanization. For example, urban hedgehogs alter their foraging behaviour to avoid crowded areas in daylight (Dowding et al., 2010), great tits use higher pitch in their calls in the noisy urban environment (Slabbekoorn and Peet, 2003; Zollinger et al., 2017), etc. On the contrary, the direct impact of human behaviour on animals has mostly been discussed using pet animals (Hosey and Melfi, 2014) and studies pertaining to the direct human impact on free-ranging animals are lacking. Free-ranging dogs (*Canis lupus familiaris*) are an excellent model system to evaluate the impact of interspecific interactions with humans in ecologically relevant contexts. These dogs regularly interact with humans in all possible human habitations in most of the developing countries (Sen Majumder et al., 2014; Vanak and Gompper, 2009). They substantially differ from pet dogs in terms of human socialisation, which in turn affects their learning ability (Brubaker et al., 2017; Brubaker et al., 2019, *in press*). Learning further allows individuals to fine-tune their behaviour to local environmental conditions by incorporating behavioural plasticity (Galef, 1995; Komers, 1997; Mery and Burns, 2010; Sol et al., 2013). Unfortunately, a limited number of studies so far have explored free-ranging dogs’ socio-cognitive dynamics and their direct interactions with humans.

Social organization in free-ranging dogs can vary from solitary to groups (sometimes up to 15 individuals, personal observation). Such flexibility in group size might have evolved as a by-product of foraging ecology and competition, but the underlying dynamics at the population level are yet to be understood. Foraging associations in free-ranging dogs are dynamic and can vary over different seasons, primarily driven by social needs (Sen Majumder et al., 2014). The social groups show interesting cooperation-conflict dynamics, with the presence of alloparenting by both related females and males on the one hand and mother-offspring conflict and milk theft on the other (Paul et al., 2014; Paul et al., 2015). Though the dogs live in stable social groups, unlike their closest living ancestors, the grey wolves (*Canis lupus lupus*), they do not display strict reproductive hierarchies and rarely hunt (Cafazzo et al., 2010; Font, 1987; Fox et al., 1975). Therefore, the evolution of flexible group size in dogs and the advantages of group living needs critical assessment. Comparative studies using individual and group level responses in various contexts can help shed light on the adaptive advantages of group living in dogs.

In an earlier study, we compared solitary individuals and groups of free-ranging dogs in problem-solving conditions (physical cognitive tasks) to understand their cognitive abilities, cooperation, and social tolerance. In spite of limited success rates in both the solitary and group conditions, cooperative motivations in terms of co-feeding and social tolerance were observed in groups (Bhattacharjee et al., under review). While such processes (social tolerance and cooperation) can facilitate group living, a more robust understanding of their adaptability to human habitats can be developed by observing their direct interactions with humans, focusing largely on their ecology. Free-ranging dogs have earlier been shown to comprehend context-dependent (friendly, threatening, etc.) human social cues. Their situation-specific responses to such cues reflect a great deal of understanding of human intentions, which is also vital for their survival in the human-dominated environment (Bhattacharjee et al., 2018). In this study, we aim to understand (a) the effects of varying human social cues on groups of free-ranging dogs, (b) differences in group and individual level responses (comparative approach), and (c) intra-group behavioural differences of individuals.

Living in groups sometimes help members to react or respond to a cue (stimuli) differently from a solitary individual. For example, a threatening signal can impact a solitary individual with a greater magnitude as compared to a group of individuals, where the impact of the threat would be reduced to a significant extent because of a ‘dilution effect’ (Lima, 1995; Stankowich and Blumstein, 2005). However, intra-group differences among individuals can still be present and get reflected in group responses. A major contributing factor responsible for differential outcomes to the same cue can be dominance-rank relationships within social groups (Francis, 2010; Rowell, 1974). Unfortunately, no studies till date have examined the relationship between personality and dominance in free-ranging dogs. This study is the first attempt to gather baseline information on group-level behavioural reactions to human social cues.

In India, free-ranging dogs are often considered as a menace and consequently beaten, shooed away, and even killed (Paul et al., 2016). Although they depend heavily on humans for sustenance, avoidance of direct contact with unfamiliar humans is also observed in free-ranging dogs, but social facilitation from humans can help dogs build trust with strangers (Bhattacharjee et al., 2017b). These dogs have also been shown to adjust their point-following behaviour flexibly based on the reliability of humans (Bhattacharjee et al., 2017a). Hence, it is evident from the prior studies that these dogs have a broad behavioural repertoire that allows them to behave flexibly, adjusting their responses to humans in a situation-specific manner. We hypothesize that groups of dogs would react to the different human social cues in a similar situation-specific manner. We used published data on solitary dogs’ responses to human social cues from Bhattacharjee et al., 2018 for comparative analysis with the group-level data We also hypothesize that groups would display less anxious behavioural reactions to threatening cues as compared to solitary individuals as a result of the dilution effect. Additionally, intra-group behavioural differences would be present due to variations in personality traits. We expect no effect of sex as a function of inter-individual differences in the reactions towards the social cues.

## Methods

### A. Subjects and study areas

We tested 80 adult-only groups of free-ranging dogs with a minimum group size of 3 (average group size: 3.53 ± 0.89). Individuals that were sighted to be either resting or moving together, up to a distance of 1 meter of each other, were considered as a group. Groups were located randomly in the following areas - Kalyani (22°58’30”N, 88°26’04”E), Kolkata (22°57’26”N, 88°36’39”E), Mohanpur (22°56’49”N and 88°32’4”E) and Sodepur (22°69’82”N and 88°38’95”E), West Bengal, India. No prior information regarding the composition and location of the groups tested were available. Sexes of all the dogs were determined by observing their genitalia and additionally, phenotypic details such as coat colour, scar marks were recorded to prevent resampling. To further rule out any possibility of resampling, we tested groups from different locations on different days.

### B. Experimental Procedure

We used three different types of social cues to investigate the response of free-ranging dog groups towards an unfamiliar human. Each group was tested only once with a randomly assigned cue. An additional set of 20 groups was tested without any cue and were considered as the control set. The experimental procedure detailed below is identical to the one followed earlier for solitary dogs (Bhattacharjee et al., 2018), and is reported here again for convenience. Experimentation was carried out wherever the groups were found (e.g., streets, markets, residential areas, etc.). Thus, it can be assumed that all groups were tested within their territories. Two experimenters, namely E1 and E2, were involved and consistent throughout the study. Both E1 and E2 were young males, 28 years old, 160 – 165 cm in height with a similar physical build. The videos were recorded using a Sony HDR-PJ410 camera mounted on a tripod.

i. **Attention seeking phase -** E2 attracted the attention of a group of dogs using vocalisations for 1-2 seconds (Bhattacharjee et al., 2017a).
ii. **Transition phase -** Once the dogs were alerted, E2 left the place and stood behind the camera. E2 made sure that all the members of a group were informed. E1 appeared at the position where E2 was standing initially. The duration of this phase was 10 seconds.
iii. **Social cue phase (SCP) –** E1 stood approximately 1.5 meters away from the dogs, facing them. E1 had to adjust his position to maintain the approximate distance of 1.5 meters (since dogs were not on a leash). Upon standing, E1 provided any of the following social cues for 30 seconds, and 20 groups were tested with each of the cues detailed below.
  - *Friendly Cue (FC) -* E1 enacted a friendly gesture by bending slightly forward, extending both his arms. E1 gazed towards the dogs while providing the cue, but refrained from touching (in case of approach) the dogs deliberately to avoid any potential contact bias.
  - *Low impact threatening (LIT) -* E1 raised one of his hands (counterbalanced), kept it motionless and gazed at the dogs. This cue was used to emulate a low level of threat that people often use to shoo away dogs.
  - *High impact threatening (HIT) -* E1 used a 0.45-meter long wooden stick in his hand (counterbalanced) to provide an enhanced version of the LIT cue. E1 was facing the dogs while enacting the gesture (see Supplementary **Movie 1**). The HIT cue was considered to be a more severe threat than LIT and is also a typical behaviour observed in Indian streets.
  - *Neutral Cue (NC) / Control -* E1 stood in a neutral posture, looked straight ahead without providing any cue.
iv. **Food transfer phase -** E2 arrived and handed over a piece of raw chicken (food reward) to E1 and left. Food transfer was carried out quickly (≤ 10 seconds) without allowing the dogs to see it.
v. **Food provisioning phase (FPP) –** E1 placed the food reward on the ground, approximately 0.3-meter in front of him, thus at a distance of ∼ 1.2 meters from the dogs. E1 did not make any eye contact with the dogs after placing the food reward. FPP was carried out for 30 seconds or until a dog (or dogs) obtained the food, whichever was earlier.

### C. Data Analysis and statistics

We coded the following parameters – approach and no approach (SCP and FPP), first reaction (SCP), human proximity (SCP), latency (FPP), duration of gazing (SCP and FPP), and duration of feeding time (FPP) (see **Table S1**). A particular behavioural outcome was treated as a group response when the majority of the group members exhibited it (for numerical data, the average value was taken). During data analyses, we paid attention to both group-level responses and intra-group behavioural differences. First, we quantified the parameters mentioned above to find out free-ranging dog groups’ understanding of human social cues, and then we compared the group responses with solitary dogs’ behavioural outcomes using the earlier data. We built an index called the ‘Response Index’ (RI) to better understand free-ranging dogs’ responsiveness to human social cues when present solitarily and in groups (**Table 1**). RI included the following factors - latency to approach, the position of an individual, feeding in human proximity, and gazing at E1 and conspecifics. RI had a scale of 4 – 15, which was further divided into three categories – “High Response” (scores: 12 – 15), “Intermediate Response” (scores: 8 – 11), and “Low Response” (scores: 4 – 7). Higher RI values were considered to be indicative of dogs’ ‘sociability’ and ‘bold’ behavioural tendencies, while lower values suggested a ‘fearful’ and ‘shy’ behavioural repertoire. Although RI had the capacity of measuring intra-group differences, it could not assess the personality traits (or temperament) due to a lack of test repeatability (in various contexts) in the given experimental set-up.

**Table 1.** Response index incorporating the parameters and their corresponding scores.

| **1. Latency to approach** |  |
| --- | --- |
| **Category** | **Score** |
| 1 – 2 seconds | 4 |
| 3 – 5 seconds | 3 |
| 6 – 9 seconds | 2 |
| > 10 seconds | 1 |
| No latency | 0 |
| **2. Position of an individual** |  |
| **Category** | **Score** |
| Approach | 3 |
| Same | 2 |
| Distant | 1 |
| **3. Feeding in human proximity** |  |
| **Category** | **Score** |
| Yes | 2 |
| No | 1 |
| **4. Gazing at E1** |  |
| **Category** | **Score** |
| No | 3 |
| Short (1 – 2 seconds) | 2 |
| Prolonged ( > 3 seconds) | 1 |
| **5. Gazing at conspecifics** |  |
| **Category** | **Score** |
| No | 3 |
| Short (1 – 2 seconds) | 2 |
| Prolong ( > 3 seconds) | 1 |

We used non-parametric tests throughout the analyses. Generalized linear mixed model (GLMM) analysis was carried out using “lme4” package of R Studio. A naïve coder coded 20% of the data to check inter-rater reliability, and it was found to be very high (Approach: Cohen’s kappa = 1.00; Proximity: Cohen’s kappa = 0.85; Gazing: Cohen’s kappa = 0.86; Latency: Cohen’s kappa = 0.88). The alpha level was 0.05, but was adjusted using Bonferroni correction for post-hoc comparisons, whenever required. We coded all the behaviours from the videos in a frame-by-frame manner using Pot player (version 1.7.18344). Statistical analyses were performed using R (R Development Core Team, 2015) and StatistiXL version 1.11.0.0.

## Results

### A. Group-level response

#### Approach

In SCP, 12 groups out of 20 approached even when they received no cue (NC). Later, the number increased to 17 in the FPP, but the two response levels were not significantly different (χ^2^ Goodness of fit: χ^2^ =0.862, df = 1, p = 0.353). Similar to NC, we found the number of approaches between the two phases to be comparable for FC (no. of approaches – SCP - 20, FPP – 20, χ^2^ Goodness of fit: χ^2^ =0.000, df = 1, p = 1.000) and LIT (no. of approaches – SCP – 6, FPP – 13, χ^2^ Goodness of fit: χ^2^ = 2.579, df = 1, p = 0.108) conditions. We found a difference between the responses in the two phases of the HIT condition (no. of approaches – SCP – 0, FPP – 6, χ^2^ Goodness of fit: χ^2^ = 6.000, df = 1, p = 0.014, see **Fig. 1a**).

**Fig 1.**
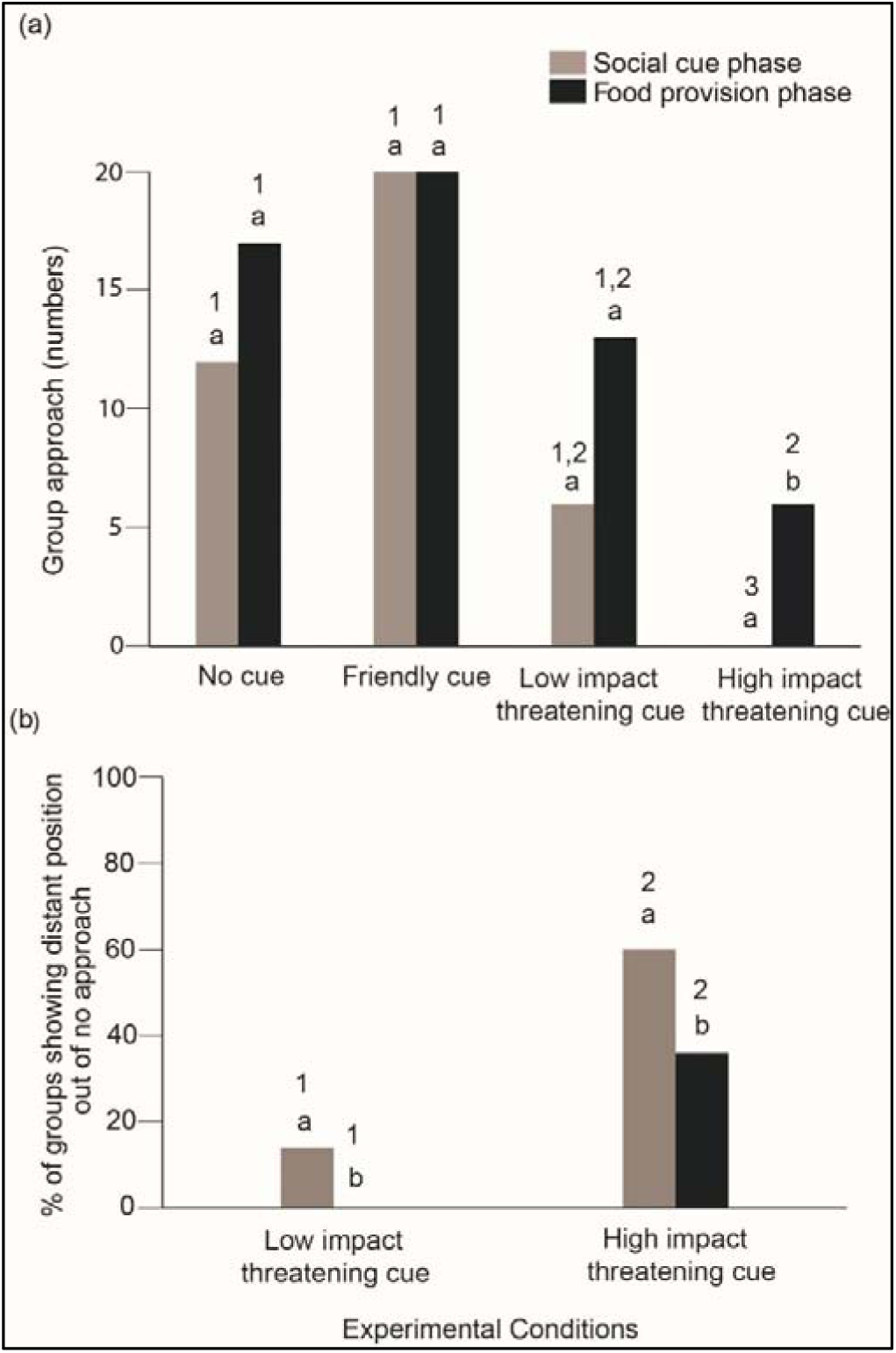
Approach and no approach. (a) Bar graph showing the number of groups showing approach responses in the two phases of the four cue conditions. (b) Bar graph showing the percentage of groups showing distant (position) no approach out of the total no approach. “a” and “b” indicate significant differences within the categories and “1” and “2” indicate significant differences between the categories.

Across conditions, we found the following results (**Fig. 1a**) - a higher number of groups approached in the SCP of FC than both LIT (χ^2^ Goodness of fit: χ^2^ = 7.538, df = 1, p = 0.006) and HIT (χ^2^ Goodness of fit: χ^2^ = 20.000, df = 1, p < 0.001) conditions. Groups were also found to approach more in the SCP of the NC than the HIT condition (χ^2^ Goodness of fit: χ^2^ = 12.000, df = 1, p = 0.001). We did not find comparisons between NC – FC, NC – LIT to be significant (**Table S2**). The FPP of HIT differed from both the NC (χ^2^ Goodness of fit: χ^2^ = 5.261, df = 1, p = 0.02) and FC (χ^2^ Goodness of fit: χ^2^ = 7.538, df = 1, p = 0.006) conditions. There was no difference in the responses between NC – FC, NC – LIT, FC – LIT, and LIT – HIT conditions of FPP.

#### No approach

We observed ‘distant no approach’ only in the LIT and HIT conditions. The differences between SCP and FPP of the two conditions were significant (Contingency Table χ^2^: χ^2^ = 7.804, df = 1, p = 0.005, **Fig. 1b**). Both in the LIT and HIT conditions, we obtained significantly higher ‘distant no approach’ in SCP compared to FPP (LIT – χ^2^ Goodness of fit: χ^2^ = 14.000, df = 1, p < 0.001; HIT – χ^2^ Goodness of fit: χ^2^ = 6.000, df = 1, p = 0.01). We also found the across-category comparisons to be significantly different (SCP - χ^2^ Goodness of fit: χ^2^ = 28.595, df = 1, p < 0.001; FPP - χ^2^ Goodness of fit: χ^2^ = 36.000, df = 1, p < 0.001). The number of ‘distant no approaches’ were significantly higher in HIT for both the phases compared to LIT.

#### First behaviour during social cue

All the groups reacted upon receiving the social cues. Gazing, gazing with tail wagging and scared were the specific responses that have been observed across conditions. In the NC condition, we found all the groups showing gazing behaviour towards E1. None of the groups showed gazing with tail wagging or scared responses (**Fig 2a**). Groups showed both gazing and gazing with tail wagging behaviours in the FC condition at equal levels (χ^2^ Goodness of fit: χ^2^ = 3.200, df = 1, p = 0.07), but did not display scared responses (**Fig 2b**). In the LIT condition, groups showed scared responses significantly more than gazing with tail wagging (χ^2^ Goodness of fit: χ^2^ = 6.231, df = 1, p = 0.01), but gazing and scared responses were comparable (χ^2^ Goodness of fit: χ^2^ = 0.889, df = 1, p = 0.34, **Fig 2c**). Gazing and gazing with tail wagging behaviours were also comparable (χ^2^ Goodness of fit: χ^2^ = 2.778, df = 1, p = 0.09). HIT condition had a strong impact on dogs as all the groups showed only scared responses (**Fig 2d**).

**Fig 2.**
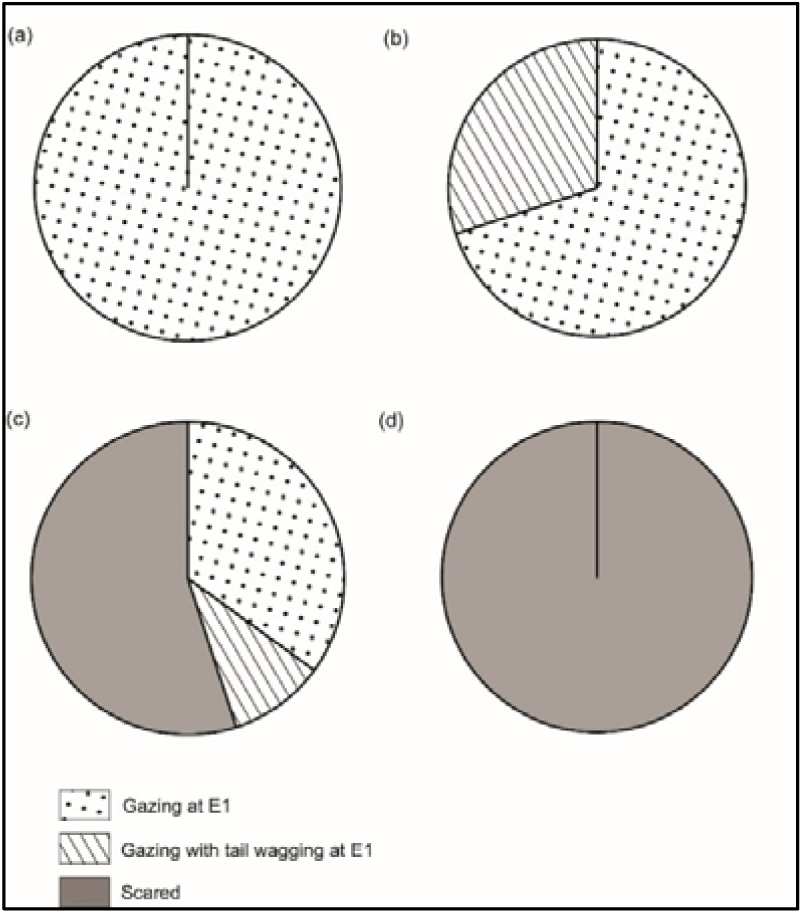
First behaviour during social cue. Pie chart showing the percentage of different behaviours during the social cues provided in (a) NC, (b) FC, (c) LIT, and (d) HIT conditions.

#### Human proximity

Groups showed variations in the duration of human proximity for different cues (Kruskal Wallis test: χ^2^ =47.259. df = 3, p < 0.001). Post-hoc pairwise comparisons revealed that groups spent a significantly higher amount of time near E1 in the FC, as compared to the NC, LIT, and HIT conditions (**Table S2**). We also found a significantly higher duration of proximity to E1 in the NC compared to the HIT condition (**Table S2**). However, we did not obtain any difference between the NC - LIT, and LIT – HIT conditions (**Table S2**).

#### Gazing

Generalised linear mixed model (GLMM) analysis revealed significant effects of the types of cues, and SCP on the duration of gazing at E1 (**Table S3**). We also compared the cumulative (pooled for all cues) duration of gazing between SCP and FPP (Mann-Whitney U test: U = 70358.500, df1 = 283, df2 = 283, p < 0.001, **Fig 3**). Across-phase comparisons revealed higher duration of gazing in SCP for each of the cues (NC – Mann-Whitney U test: U = 4058.500, df1 = 68, df2 = 68, p < 0.001; FC - Mann-Whitney U test: U = 4549.000, df1 = 68, df2 = 68, p < 0.001; LIT - Mann-Whitney U test: U = 3030.500, df1 = 62, df2 = 62, p < 0.001; HIT - Mann-Whitney U test: U = 5827.500, df1 = 85, df2 = 85, p < 0.001).

**Fig 3.**
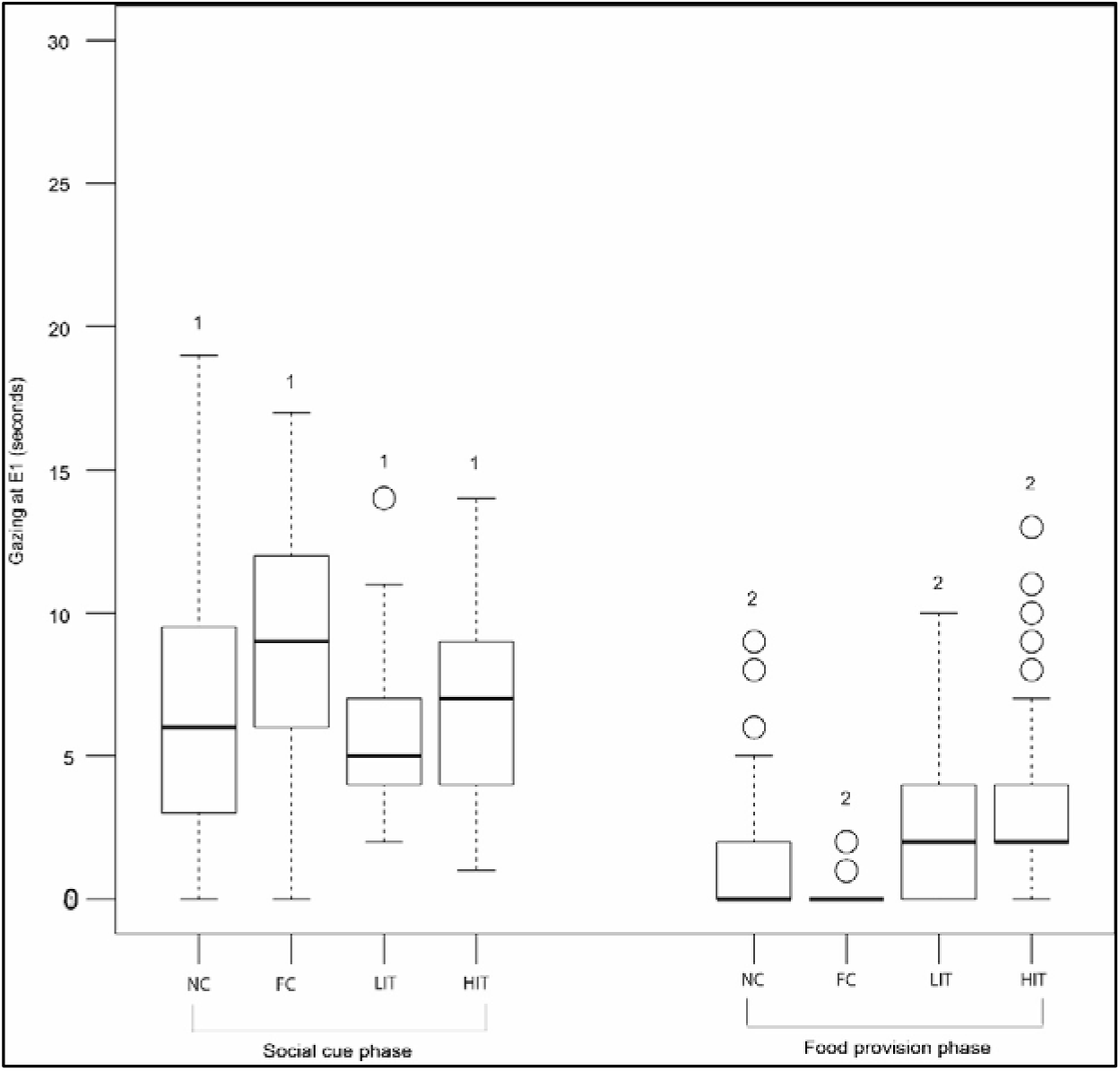
Duration of gazing at E1. Box and Whiskers plot showing the duration of gazing at the E1. Boxes represent interquartile range, horizontal bars within boxes indicate median values, and whiskers represent the upper range of the data. “1” and “2” indicate significant differences between the categories (between social cue and food provision phase).

#### Latency and duration of feeding

The latencies of the first members that approached in the FPP of the four conditions (*N* = 57) showed significant variation (Kruskal-Wallis test: χ^2^ =34.011, df = 2, p < 0.001). Dogs showed a tendency to approach significantly faster in the FC than the NC, LIT, and HIT conditions (**Table S2**). We also found differences between NC and LIT, with dogs showing faster approach in NC (**Table S2**). However, we did not see any difference between LIT and HIT, and NC – HIT conditions (**Table S2**).

The feeding time comparable among the four conditions (Kruskal-Wallis test: χ^2^ =1.161, df = 3, p = 0.762). We did not observe the group members sharing food among themselves in any of the conditions.

### B. Comparison of individual and group responses

We compared five major parameters across the two sets of experiments – approach, first behaviour after social cue, latency, proximity to human, and duration of gazing.

#### Approach

Groups showed a higher approach rate than solitary individuals (χ^2^ Goodness of fit: χ^2^ = 15.933, df = 1, p < 0.001).

#### First behaviour after social cue

The first reactions differed between the individual and group levels (**Fig 4**). Groups showed a significantly higher duration of gazing behaviour (at E1) than solitary individuals (χ^2^ Goodness of fit: χ^2^ =25.752, df = 1, p < 0.001). Apart from gazing, all the other responses were displayed at a higher rate by the solitary dogs (gazing with tail wagging - χ^2^ Goodness of fit: χ^2^ =8.526, df = 1, p = 0.004; scared - χ^2^ Goodness of fit: χ^2^ =11.792, df = 1, p = 0.001; no reaction - χ^2^ Goodness of fit: χ^2^ =8.000, df = 1, p = 0.005).

**Fig 4.**
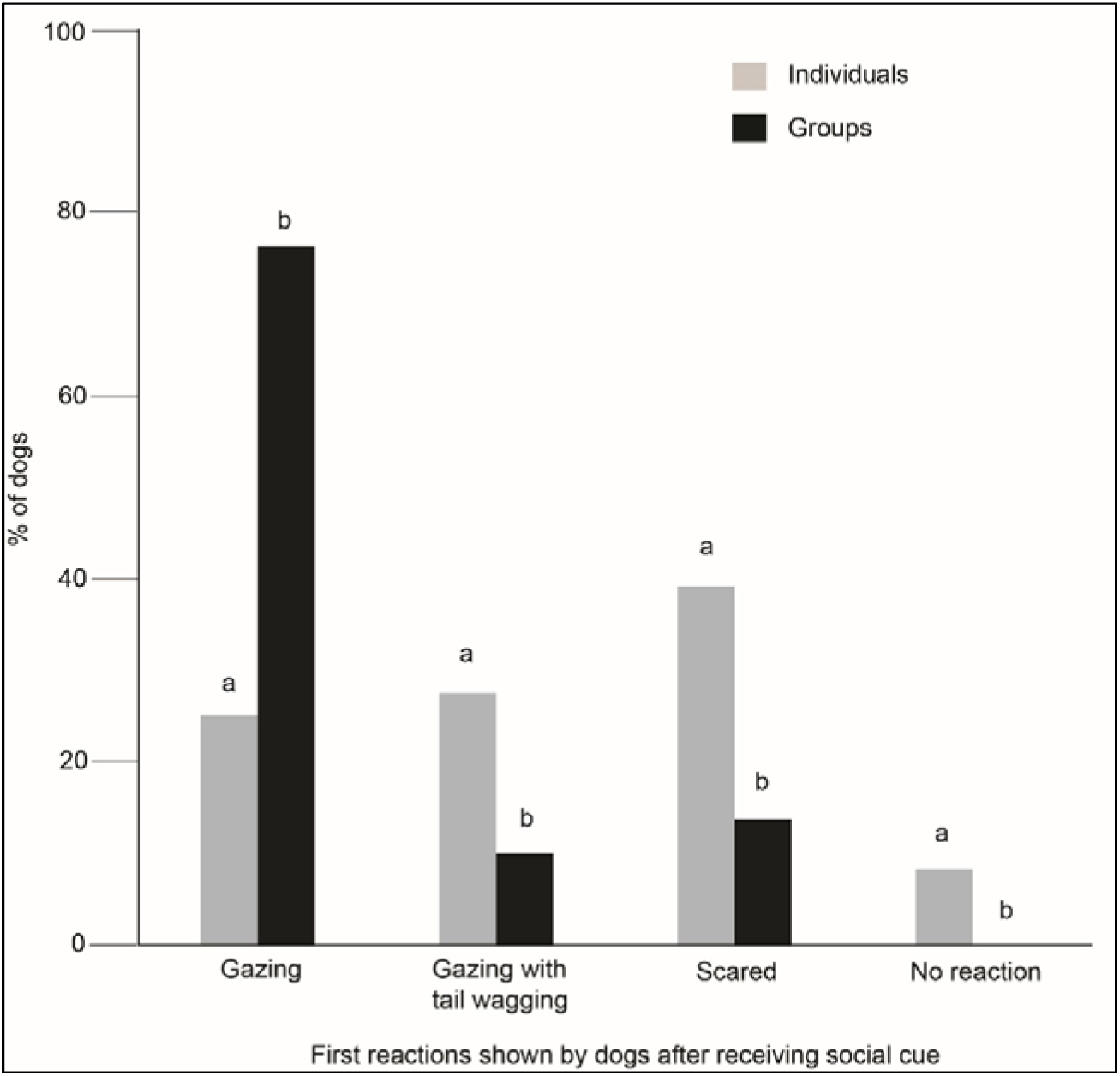
Comparison of first behaviours between solitary and groups of dogs. Bar graph showing the percentage of behaviours (first reactions in the SCP) shown by the solitary and groups of dogs towards the E1.

#### Latency

Latencies were comparable between individuals and groups for all the conditions in FPP (NC - Mann-Whitney U test: U = 177.000, df1 = 17, df2 = 17, p = 0.27; FC - Mann-Whitney U test: U = 318.500, df1 = 20, df2 = 30, p = 0.71; LIT - Mann-Whitney U test: U = 85.500, df1 = 13, df2 = 13, p = 1.00; HIT - Mann-Whitney U test: U = 4.500, df1 = 6, df2 = 1, p = 0.57).

#### Duration of proximity to E1

Generalised linear model (GLM) analysis showed significant effects of groups and solitary conditions and cue types on the duration of proximity to E1 (**Table S4**). Overall, the duration of proximity was found to be significantly higher for the groups (4.41 ± 5.97 sec) as compared to individuals (3.45 ± 7.36 sec).

#### Duration of gazing at E1

GLM analysis revealed significant effects of groups and solitary conditions, cue types, and phases on the duration of gazing at E1 (**Fig S1, Table S5**). We found both individual and interactive effects of predictors (dog composition type, cue, phase) on the gazing duration. Gazing was found to be significantly higher in SCP for both the individuals and groups, as compared to FPP (Individuals - Mann-Whitney U test: U = 10749.000, df1 = 120, df2 = 120, p < 0.001; Groups - Mann-Whitney U test: U = 5868.000, df1 = 80, df2 = 80, p < 0.001). We also found a difference between the individuals and groups in FPP (Mann-Whitney U test: U = 5831.500, df1 = 120, df2 = 80, p = 0.01) with individuals showing higher duration of gazing. However, the gazing duration was comparable in the SCP phase (Mann-Whitney U test: U = 4935.500, df1 = 120, df2 = 80, p = 0.73).

### C. Intra-group differences

#### Response Index

RI values differed between the different cues (Kruskal Wallis test: χ^2^ = 100.320, df = 3, p < 0.001). Post-hoc pairwise comparisons revealed significant differences between NC – FC (Mann-Whitney U test: U = 3037.500, df1 = 67, df2 = 68, p = 0.001, higher RI values in FC), NC – HIT (Mann-Whitney U test: U = 4583.000, df1 = 68, df2 = 85, p < 0.001, higher RI values in NC), FC – LIT (Mann-Whitney U test: U = 3292.500, df1 = 68, df2 = 62, p < 0.001, higher RI values in FC), FC – HIT (Mann-Whitney U test: U = 5357.000, df1 = 68, df2 = 85, p = 0.73, higher RI values in FC), and LIT – HIT (Mann-Whitney U test: U = 3871.500, df1 = 62, df2 = 85, p < 0.001, higher RI values in LIT). We did not find any difference between NC – LIT (Mann-Whitney U test: U = 2585.000, df1 = 68, df2 = 62, p = 0.02). Additionally, 25%, 95%, 0%, and 10% of the groups showed the highest RI value (i.e. “15”) in NC, FC, LIT, and HIT conditions respectively. We also calculated the percentages of the groups showing RI values ranging from 12 to 15 (designated as high responders). We found that 70%, 100%, 45%, and 35% of the groups obtained RI values from 12 to 15 in NC, FC, LIT, and HIT conditions.

In the NC condition, 14 groups had high responders; out of these, three groups had more than one individual as high responder (χ^2^ Goodness of fit: χ^2^ =4.571, df = 1, p = 0.03). In the FC condition, all 20 groups had one or more individuals as high responders, out of which, seven groups had only one high responder (χ^2^ Goodness of fit: χ^2^ =1.800, df = 1, p = 0.18, **Fig 5a**). We found all nine groups in the LIT condition to have only one member as high responder (χ^2^ Goodness of fit: χ^2^ =9.000, df = 1, p = 0.003, **Fig 5b**). In the HIT condition, only one of the seven groups had multiple high responders (χ^2^ Goodness of fit: χ^2^ =3.571, df = 1, p = 0.05, **Fig 5c**).

**Fig 5.**
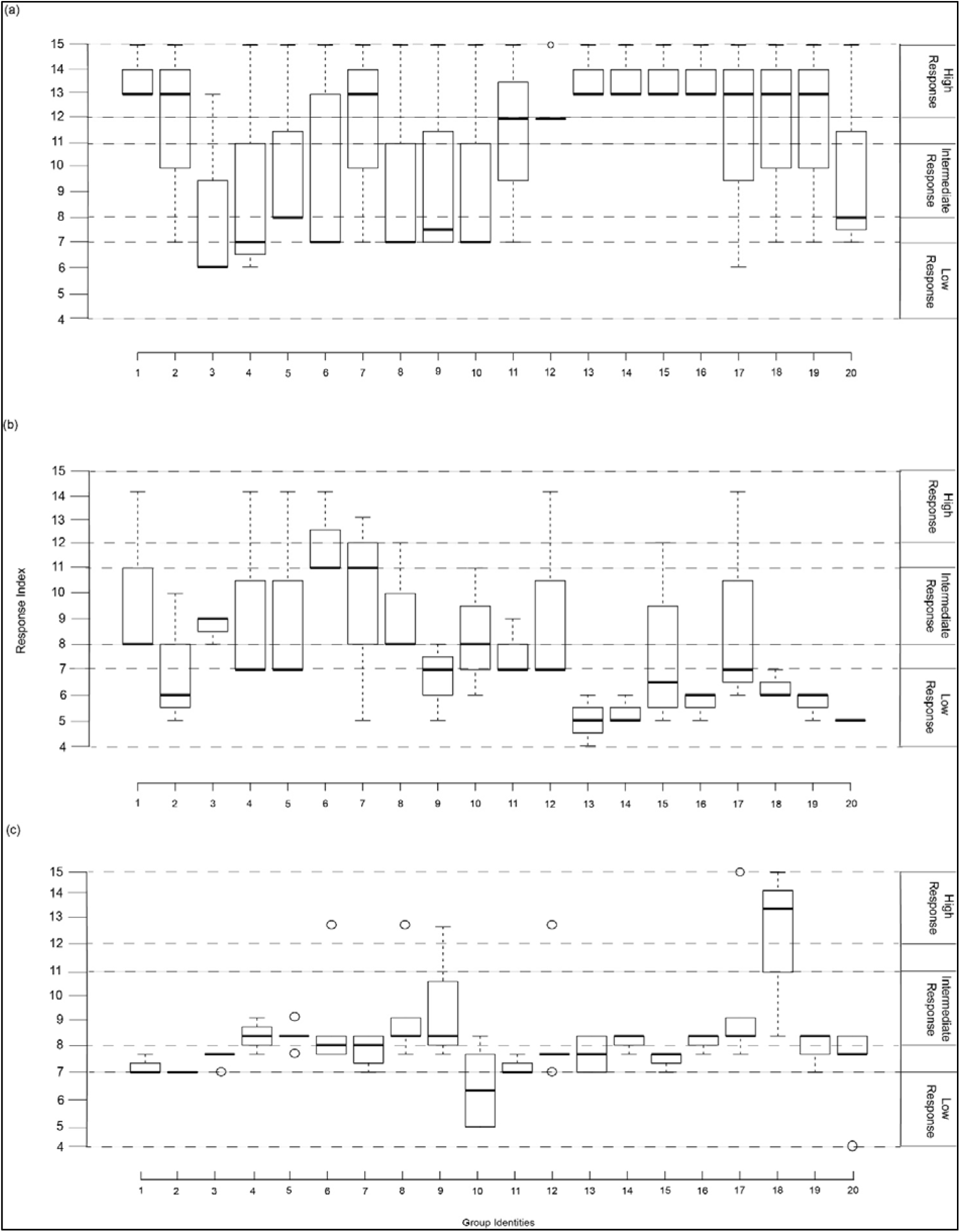
Response Index (RI) Box and Whiskers plot showing the distribution of values of the RI in (a) FC, (b) LIT, and (c) HIT conditions.

#### Effect of sex on high responders

We found that overall (all cue types, pooled data), 52 males and 30 females were high responders (χ^2^ Goodness of fit: χ^2^ =5.902, df = 1, p = 0.01). We did not find any difference at the sex ratio of the total dogs tested in the study (χ^2^ Goodness of fit: χ^2^ =2.972, df = 1, p = 0.08). We further analysed the responses in the two threatening cue conditions and found that the number of male high responders were higher than females (χ^2^ Goodness of fit: χ^2^ =8.000, df = 1, p = 0.005), suggesting that males might be bolder than females.

## Discussion

This study corroborates earlier findings of free-ranging dogs’ situation-specific responses towards varying human social cues (Bhattacharjee et al., 2018). Our results highlight the differences between solitary and group-level reactions, with dogs showing a “bolder” response when in groups. We further provide the first evidence of sex difference in the bold behavioural tendency of free-ranging dogs while responding to threatening cues from humans. Higher approach rates, less anxious or fearful behaviours were the key features that differentiated the response of dog groups from that of the solitary individuals to threatening cues, suggesting a direct benefit of group-living over a solitary lifestyle.

The general pattern of response to the different cues by groups was similar to that of the solitary dogs. However, the approach rate was found to be higher in groups, especially in the SCP of LIT, providing evidence of a less effective LIT cue when the dogs were in a group. In India, solitary dogs on streets are more prone to receive threatening signals from humans as compared to groups of dogs (personal observations). It could also be a consequence of the higher perception of threat or shyness towards unfamiliar humans that solitary dogs avoid making direct physical contact with unfamiliar humans (Bhattacharjee et al., 2017b). Studies show that animals living in groups are less vigilant than their solitary counterparts in various ecological contexts (Delm, 1990; Dimond and Lazarus, 1974; Quenette and Gerard, 1992). However, in our experiments, gazing at the experimenter as a reaction to social cues was found to be a significant behaviour in the free-ranging dog groups. We suspect that the free-ranging dogs, due to the constant anthropogenic stress in their environment, are naturally vigilant, and the gazing response is a part of their behavioural repertoire. Moreover, they are territorial and need to defend their territories from intruders, including humans, other dogs, and other animals, giving rise to a complex and dynamic behavioural system. They typically defend their territories in groups, while solitary dogs typically are more prone to avoid situations of conflict either with other dogs or humans.

Our results revealed an interesting pattern regarding the behavioural tendencies of groups. At the intra-group level, dogs differed in terms of their responses, e.g. a majority of the dogs were high responders in the FC condition. Though there was a gradual decrease in the number of high responders from FC to the threatening cue conditions (LIT and HIT), they nevertheless were not absent in the situations of threat. This suggests that within a group, there are individuals with varying personalities/ temperaments, and the high responders can be considered to show “bold” behavioural tendencies. It should be noted that males tended to be bolder than females, in this context. This study opens up the need for further explorations into context-dependent responses in free-ranging dog groups to understand how different behavioural types emerge in the groups and the underlying role of sex in the development of a bold temperament.

Free-ranging dogs, irrespective of their social composition enact situation-relevant reactions to commonly used human social cues. Our results suggest a potential advantage of group living in dogs over a solitary lifestyle when it comes to interacting with humans, especially in unfavourable circumstances. This ecological advantage need not be driven by the benefits of kin selection (Hamilton, 1964), but would nevertheless be amplified in the evolutionary timescale, if group members are closely related to each other, which often tends to be the case (Paul et al., 2015). While a certain degree of difference is evident, solitary and groups of free-ranging dogs mostly overlapped in their pattern of responses, probably depicting the best possible strategy adapted to living in the human-dominated environment. Therefore, we assume that a lack in supply of ample amounts of human subsidized food and competition could be the potential conflicting factors that ultimately influence group size and stability, causing a flexible nature of social composition in free-ranging dogs. Future studies using the postulates of ‘Resource Dispersion Hypothesis’ (RDH) would be useful to have vital information on the mechanisms that govern group formation and splitting in dogs (Macdonald and Johnson, 2015). Information regarding the potential differences between the behavioural tendencies of free-ranging dogs can further be checked by linking dominance-rank relationships.

Our study revealed significant insights into the dog-human relationship on the streets. Understanding the intents of humans is crucial for these dogs to adjust their behavioural responses accordingly. Above all, these situation-relevant responses to human social cues can provide us with the direction required for tackling and mitigating the rapidly increasing free-ranging dog-human conflict in most of the developing countries.

## Acknowledgements

DB would like to thank DST INSPIRE for providing the research fellowship. The authors thank IISER Kolkata for infrastructural support.

## Funding statement

We received no funding for the study.

## Authors’ contributions

Conceptualization: DB and AB; Methodology: DB and AB; Investigation: DB and SS; Analysis: DB; Original Draft: DB; Review and Editing: DB and AB; Resources: AB; Supervision: AB.

## Competing interests

Authors declare no competing interests.

